# Topographic organization of the “third tier” dorsomedial visual cortex in the macaque

**DOI:** 10.1101/519009

**Authors:** Kostas Hadjidimitrakis, Sophia Bakola, Tristan A. Chaplin, Hsin-Hao Yu, Omar Alanazi, Jonathan M. Chan, Katrina H. Worthy, Marcello G.P. Rosa

**Affiliations:** Department of Physiology, Biomedicine Discovery Institute, Monash University, Clayton, VIC 3800, Australia.; ARC Centre of Excellence for Integrative Brain Function, Monash University Node, Monash University, Clayton, VIC 3800, Australia.; School of Biomedical Sciences (previously Vision, Touch and Hearing Research Centre, Department of Physiology and Pharmacology), The University of Queensland, Brisbane, QLD 4072, Australia.

## Abstract

The boundaries of the visual areas located anterior to V2 in the dorsomedial region of the macaque cortex remain contentious. This region is usually conceptualized as including two functional subdivisions: the dorsal component of area V3 (V3d), laterally, and another area, named the parietooccipital area (PO) or V6, medially. However, the nature of the putative border between V3d and PO/V6 has remained undefined. We recorded the receptive fields of multiunit clusters in adult male macaques, and reconstructed the locations of recording sites using histological sections and “unfolded” cortical maps. Immediately adjacent to dorsomedial V2 we observed a representation of the lower contralateral quadrant, which represented the vertical meridian at its rostral border. This region, corresponding to V3d of previous studies, formed a simple eccentricity gradient, from approximately <5° in the annectant gyrus, to >60° in the parietooccipital sulcus. However, there was no topographic reversal where one would expect to find the border between V3d and PO/V6. Rather, near the midline, this lower quadrant map continued directly into a representation of the peripheral upper visual field, without an intervening lower quadrant representation that could be unambiguously assigned to PO/V6. Thus, V3d and PO/V6 form a continuous topographic map, which includes parts of both quadrants. Together with previous observations that V3d and PO/V6 are both densely myelinated relative to adjacent cortex, and share similar input from V1, these results suggest that they are parts of a single area, which is distinct from the one forming the ventral component of the third tier complex.

**Significance statement:** The primate visual cortex has a large number of areas. Knowing the extent of each visual area, and how they can be distinguished from each other, are essential for the interpretation of experiments aimed at understanding visual processing. Currently, there are conflicting models of the organization of the dorsomedial visual cortex rostral to area V2 (one of the earliest stages of cortical processing of vision). By conducting large-scale electrophysiological recordings, we found that what were originally thought to be distinct areas in this region (dorsal V3, and the parietooccipital area [PO/V6]), together form a single map the visual field. These results will help guide future functional studies, and the interpretation of the outcomes of lesions involving the dorsal visual cortex.

## Introduction

Despite four decades of research, there is still controversy regarding the boundaries of visual areas in the “third tier” cortex, i.e., the areas located rostral to the second visual area (V2, (Allman and Kaas, 1975). Here we focus on the parts of this complex located in the dorsomedial region of the macaque brain, including the medial part of the lunate sulcus, annectant gyrus, parietooccipital sulcus and mesial surface of the brain (Fig. 1). The traditional view has been that there are at least two subdivisions in this region. Laterally, in the lunate sulcus and annectant gyrus, most studies indicate the existence of a dorsal component of area V3 (V3d; Gattass et al., 1988). Medially, the cortex along the banks of the parietooccipital sulcus and mesial surface is usually assigned to a different region, termed the parietooccipital area (PO; Colby et al., 1988) or V6 (Galletti et al., 1999) (Fig. 1). Although proposals regarding the boundaries of PO/V6 continue to evolve (Gamberini et al., 2015), it is generally agreed that receptive fields of neurons in this region emphasize peripheral vision. Moreover, models proposed by various groups converge on the idea that PO/V6 encompasses a representation of the lower quadrant in the parietooccipital sulcus, as well as a representation of the upper visual field on the mesial surface of the hemisphere. Models proposed for the organization of PO/V6 in capuchin monkeys (Neuenschwander et al., 1994) and humans (Pitzalis et al., 2006) reflect these features.

**Figure 1:**
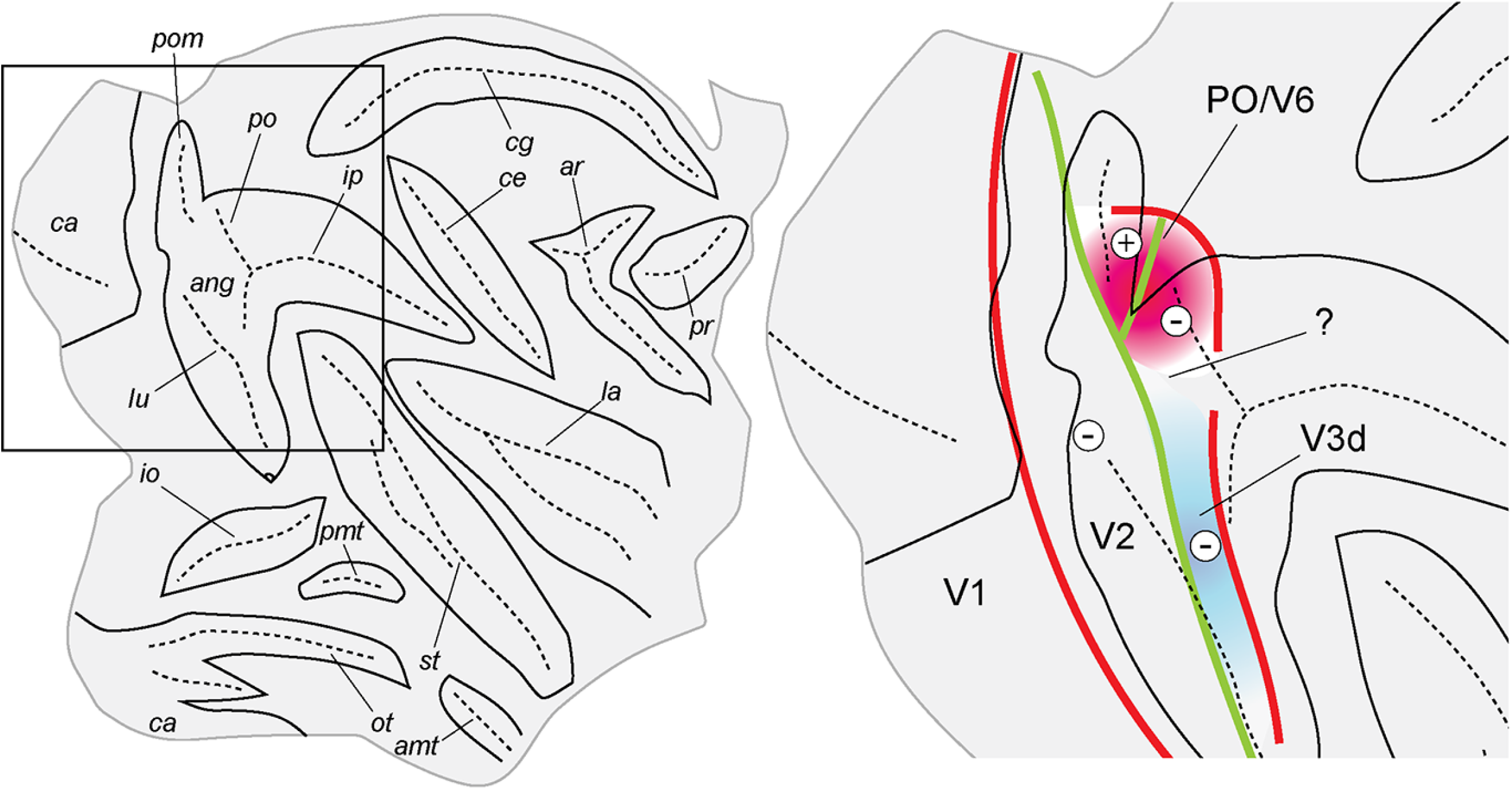
**Left**: unfolded representation of the right hemisphere of a macaque brain, based on a reconstruction prepared with CARET (Van Essen et al. 2001). In this map the lips of the sulci are indicated in continuous black line, and their main internal folds by dashed lines. The rectangle shows the region magnified on the right. **Right**: diagrammatic representation of the model we test in this paper (Ungerleider and Desimone 1986; Colby et al. 1988; Gattass et al. 1988; Gamberini et al. 2015), according to which the cortex rostral to dorsomedial area V2 contains 2 areas, V3d (blue) and PO (or V6, pink), each with a distinct representation of the lower visual field. The question mark indicates the possibility that another area, PIP, may separate V3d from PO/V6. In this diagram thick red lines indicate representations of the vertical meridian of the visual field, thick green lines indicate representations of the horizontal meridian, the – signs indicate representations of the lower contralateral quadrant, and the + signs indicate a representation of the upper contralateral quadrant. Abbreviations (sulci): amt: anterior middle temporal sulcus; ang: annectant gyrus; ar: arcuate sulcus; ca: calcarine sulcus; ce: central sulcus; cg: cingulate sulcus; io: inferior occipital sulcus; ip: intraparietal sulcus; la: lateral sulcus; lu: lunate sulcus; ot: occipitotemporal sulcus; pmt: posterior middle temporal sulcus; po: parietooccipital sulcus; pom: parietooccipital medial sulcus; pr: principal sulcus; st: superior temporal sulcus.

Visual areas corresponding to early stages of visual processing each form a representation of the visual field (e.g., Van Essen and Zeki, 1978; Sereno et al., 1995; Rosa et al., 1997). Thus, if V3d and PO/V6 are distinct areas, it would be expected that they form separable visuotopic maps. In particular, as depicted in most studies, there should be distinct representations of the peripheral lower quadrant near their common boundary (Fig. 1). However, the transition between V3d and PO/V6 has never been documented in detail, leaving significant room for other interpretations. For example, based on analyses of cortical architectonics, Lewis and Van Essen (2000) indicated that V3d and PO/V6a may not be adjacent, being separated by a posterior intraparietal area (PIP). Conversely, based on meta-analysis of published data, Angelucci and Rosa (2015) proposed that V3d and PO/V6 could actually be parts of the same area. Indeed, V3d and PO/V6 share significant similarities, such as dense myelination, a projection from the primary visual cortex (V1) that originates in layer 4b, and large numbers of direction-selective neurons (Felleman and Van Essen, 1987; Felleman et al., 1997; Galletti et al., 2001; Rosa et al., 2009; Pitzalis et al., 2010). They are also reported to have complementary emphases in representation of central (V3d) versus peripheral (PO/V6) visual field. Given the above, it is important to explore the visuotopic organization of the cortex around their putative border in order to ascertain if the 2-areas scheme outlined above does in fact provide the best description of the organization of the dorsomedial third tier cortex. Bringing clarity to the topographic organization of this region of the macaque cortex is also relevant in order to understand similarities and differences with New World monkeys, in which a single dorsomedial area (DM) has been proposed to occupy the corresponding region (Rosa and Schmid, 1995; Rosa et al., 2005).

Based on electrophysiological recordings, we found that the cortex anterior to V2 formed a relatively simple pattern of representation of the lower quadrant, with lateral to medial gradients of increasing eccentricity and receptive field size. However, we also found that this representation merged directly into a representation of the upper quadrant near the midline, thereby forming a single representation of the peripheral visual field. Complementing the results of a recent study of the lateral part of the third tier complex (Zhu and Vanduffel, 2018), these findings point to a reinterpretation of the areas rostral to dorsal V2 in the macaque.

## Materials and methods

The present report is based on data obtained in 4 young adult male macaque monkeys *(Macaca fascicularis).* The experimental protocols were approved by the Animal Experimentation Ethics Committees of the University of Queensland and Monash University, which also monitored the welfare of the animals. All procedures complied with the guidelines of the Australian Code of Practice for the Care and Use of Animals for Scientific Purposes.

### Preparation

The animals were pre-medicated with i.m. injections of diazepam (3 mg/kg) and atropine (0.2 mg/kg), and, after 30 minutes, were anesthetized with ketamine (50 mg/kg) and xylazine (3 mg/kg; cases 1-3), or with a ketamine/Dormitor/Butorphenol cocktail (0.1 mg/kg, i.m.; case 4). Anesthesia was maintained with additional doses of ketamine (12 mg/kg; cases 1-3) or alfaxan (8 mg/kg; case 4) throughout surgery. The animals were placed in a stereotaxic frame and, prior to the beginning of recording sessions, were implanted with a bolt for holding the head while allowing an unobstructed field of vision (e.g. Bourne and Rosa 2003). The femoral vein was cannulated, and a craniotomy was performed to expose the dorsal aspect of the occipital and parietal lobes. After the surgical procedures were completed, the animals were administered an i.v. infusion of pancuronium bromide (0.5 mg/kg, followed by 0.1 mg/kg/h), combined with sufentanil citrate (6–8 μg/kg/h), in a solution of sodium chloride (0.18%) / glucose (4%) and dexamethasone (0.4 mg/kg/h). They were maintained under artificial ventilation, with a gaseous mixture of N_2_O/O_2_ (7:3). Vital signs (electrocardiograph, PO2, and levels of spontaneous activity in the cortex) were continuously monitored, and the temperature was maintained between 36.5°C and 37°C by means of a heating blanket connected to a rectal temperature probe. Mydriasis and cycloplegia were induced by topical applications of atropine (1%) and phenylephrine hydrochloride (10%) eye drops. Appropriate focus was achieved by means of contact lenses with 3 mm artificial pupils, which brought into focus the surface of a 57.3 cm radius translucent hemispheric screen centered on the eye contralateral to the cerebral hemisphere to be studied.

### Electrophysiology

Tungsten microelectrodes (~1MΩ) were inserted in anteroposterior rows along parasagittal planes approximately 1.5 mm apart. Penetrations extended in most cases to the calcarine sulcus, and in many cases to the ventral surface of the cortex, including recordings from portions of V1 and V2 in these regions. Recording sites were obtained at different depths in each penetration (every 300–500 μm). Amplification and filtering were achieved via an AM Systems model 1800 microelectrode alternating current amplifier.

Visual stimuli were monocularly presented under mesopic adaptation levels to the eye contralateral to the cortical hemisphere from which the neuronal recordings were obtained. The eye ipsilateral to the recording hemisphere was occluded. Visual stimulation was achieved through a manually operated light source, which was projected onto the translucent hemisphere (Yu and Rosa, 2010). At each recording site, receptive fields of single units or small unit clusters were mapped by correlating changes in neural activity with stimulation of specific portions of the visual field. Typical visual stimuli were white spots (1 –10° in diameter) and luminous bars (2–20° long, 0.2–1° wide), moved or flashed (1 Hz) on the surface of the screen. The stimulus luminance was between 1 and 10 cd/m. Receptive fields were drawn as rectangles parallel to the axis of best orientation. The position of each electrode penetration was marked on digital images of the pattern of blood vessels on the cortical surface, obtained with a CCD camera. Electrolytic lesions (4–5 μA for 10 seconds) were placed during the experiment to mark the transitions between areas, the end of electrode tracks, and sites of special interest.

### Histology

At the end of the experiments, the animals were administered a lethal dose of sodium pentobarbitone (100 mg/kg, i.v.), and perfused transcardially with heparinized saline or phosphate buffer, followed by 4% paraformaldehyde in 0.1M phosphate buffer (pH 7.4). The brains were removed from the skull, blocked, and cryoprotected by immersion in buffered solutions of sucrose (10–30%). Once the brains sank in the 30% sucrose solution, parasagittal sections (40 or 50 μm) were obtained using a cryostat. Adjacent series were stained for Nissl substance, cytochrome oxidase and myelin, using the Gallyas (1979) procedure (Fig. 2). All sections were coverslipped with DPX, after dehydration in ethanol and clearing with xylene.

**Figure 2:**
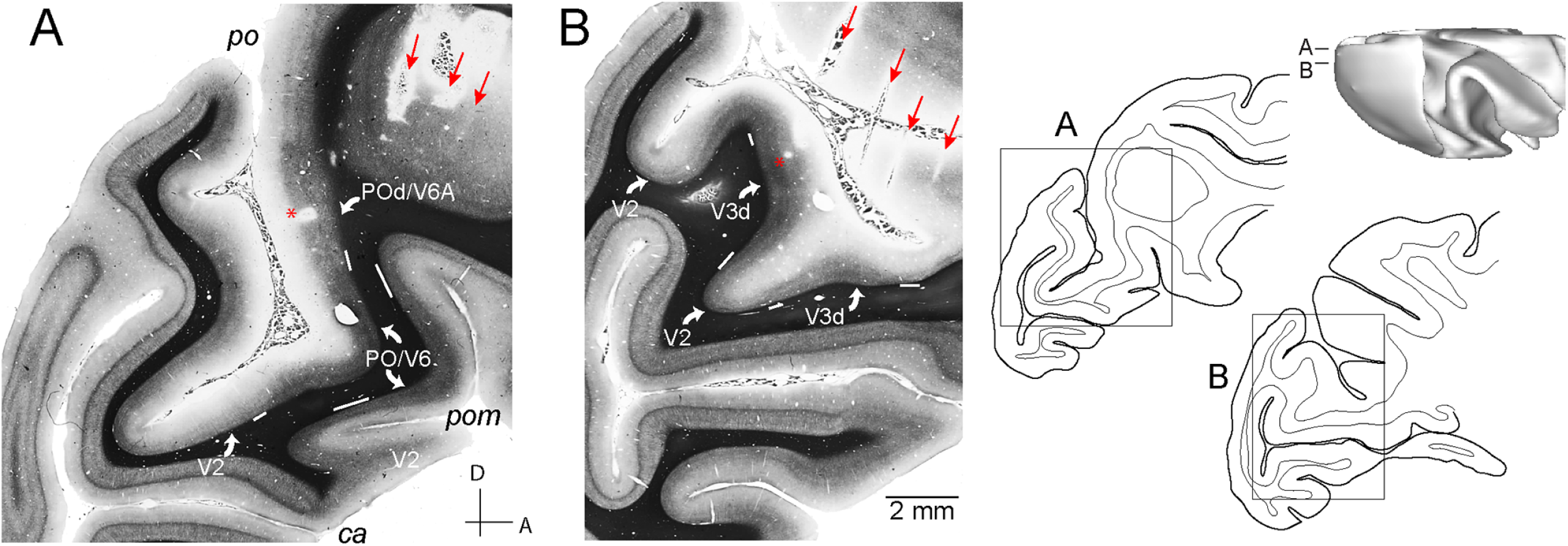
**A, B**: parasagittal sections stained with the Gallyas (1979) method, showing the histological reconstruction of the electrode penetrations and myeloarchitecture of V2, cortex rostral to V2 (here indicated as “V3d”, or PO/V6, according to previous studies), and an area in the rostral bank of the parietooccipital sulcus (POd, or V6A; Neuenschwander et al. 1994; Luppino et al. 2005). The magnified regions are indicated by rectangles in tracings of the sections (right), and their levels are indicated in a dorsal view of the brain reconstructed in CARET. Several electrode tracks (red arrows) and electrolytic lesions (asterisks) are visible, demonstrating the angle of approach of the electrodes. White lines underlying layer 6 indicate the myeloarchitectural transition zones. Both V3d and PO/V6 stand out as more densely myelinated than adjacent areas.

The positions of recording sites were reconstructed in serial sections, based on histological observation of gliosis caused by the electrode tracks (e.g. Fig. 2), electrolytic lesions, and transitions between the gray and white matter. Shrinkage due to histological processing was estimated by comparing the distances between the electrode tracks in the sections with the microdrive readings.

### Data analysis

The positions of the horizontal and vertical meridians of the visual field were estimated during the experiment by assuming that the former was aligned with the elevation of the center of the optic disk, and the latter corresponded to a vertical line approximately 15° from the center of this retinal landmark. These estimates were refined during the data analysis using well-established features of the visual topography of the macaque cortex as references: namely, we assumed that receptive fields recorded at the V1/V2 histological border overlapped with the vertical meridian of the visual field, and that receptive fields recorded from neurons near the anterior border of V2 represented the horizontal meridian (Van Essen and Zeki, 1978; Gattass et al., 1981). Repeated plotting of small receptive fields in these areas demonstrated that eye drift did not occur under the dose of pancuronium bromide that we used.

Receptive fields were digitized as rectangles on the spherical surface. The centers of the receptive fields were calculated as the mean values of the azimuth and elevation within the sectors of the visual field which they encompassed, their eccentricities as the geodesic distance between the centers of the receptive field and the intersection between the vertical and horizontal meridians, the polar angle as the angle between the centers of the receptive fields and the horizontal meridian, and the receptive field sizes as the square root of the surface areas of the rectangles that represented them (Yu and Rosa, 2010).

Three-and two-dimensional reconstructions of the cortical surface were generated with the software CARET (Van Essen et al., 2001). Sections were scanned and aligned with a graphics software (Adobe Illustrator) using pinholes created prior to sectioning as a reference. Layer 4 contours from each case were traced manually on the digitized sections, and registered. The contours were then imported into CARET to reconstruct the 3D surface models, and to create unfolded (2D) maps of the cortex. Visuotopic coordinates from each recording site were projected on the reconstructed surfaces. Subsequently, visuotopic maps were created by interpolating the eccentricity and polar angle of each recording site’s receptive field across the region of interest. We then employed an interpolation procedure to estimate coordinates at all mesh nodes, using a distance weighted smoothing algorithm (Sereno et al., 1994; Chaplin et al., 2013).

## Results

The locations of the recording sites, which yielded visual receptive fields in the region of interest, are illustrated in Figure 3. In each animal, we examined a cortical region extending approximately from the rostral border of V1 to the caudal intraparietal sulcus, and from the midline to the medial third of the lunate sulcus. The yellow lines in the unfolded maps (Fig. 3B, D, E, F) indicate the approximate medial boundary of the region encompassing recording sites that were located in the banks of the lunate, parieto-occipital and intraparietal sulci (see schematic in Fig. 3C). In each case, approximately half of the recording sites were located in the banks of these sulci. For further orientation, red asterisks also indicate the crown of the annectant gyrus (a buried gyrus, not visible from the surface of the brain, which separates the lunate sulcus from the posterior ramus of the intraparietal sulcus). The lateral limit of our recording grids included this gyrus.

**Figure 3:**
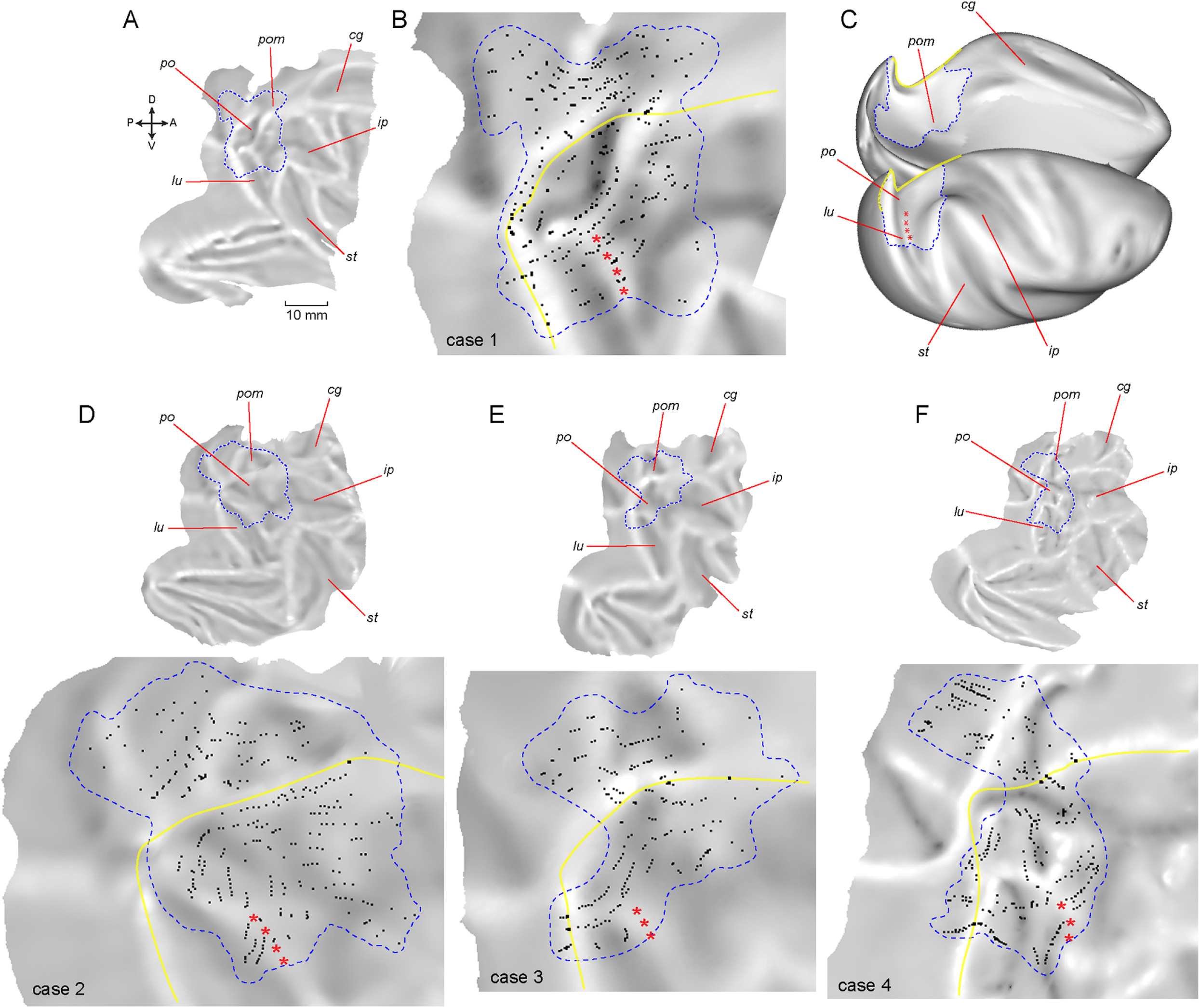
Unfolded reconstructions of the cortex in 4 cases, showing the locations of recording sites from which visual responses were obtained in dorsal extrastriate areas. **A**: reconstructed cortex in case 1, showing locations of the sulci (for abbreviations, see Fig. 1). In the insert, A, D, P and V indicate anterior, dorsal posterior and ventral in the brain. **B**: magnified view of the dorsal cortex, with recording sites indicated as black points. The dashed line indicates the outline of the region explored, used for analysis of visual topography (see Fig. 4, below). The yellow line indicates the limit between the cortex buried in the lunate-parietooccipital cleft, and the cortex exposed in the midline (including the parietooccipital medial sulcus). **C**: partially inflated view of the brain, indicating the location of the region reconstructed in B. **D-F**: similar representations of explored region and recording sites in cases 2, 3 and 4. In all maps, the crown of the annectant gyrus is indicated by the red asterisks.

Figure 4 summarizes the visual topography of the studied region in unfolded two-dimensional reconstructions. The left panels illustrate interpolated polar angle maps, and the right panels show the eccentricity maps. Receptive field sequences that illustrate in more detail the main points highlighted in the analysis below are shown in Figures 5–7.

**Figure 4:**
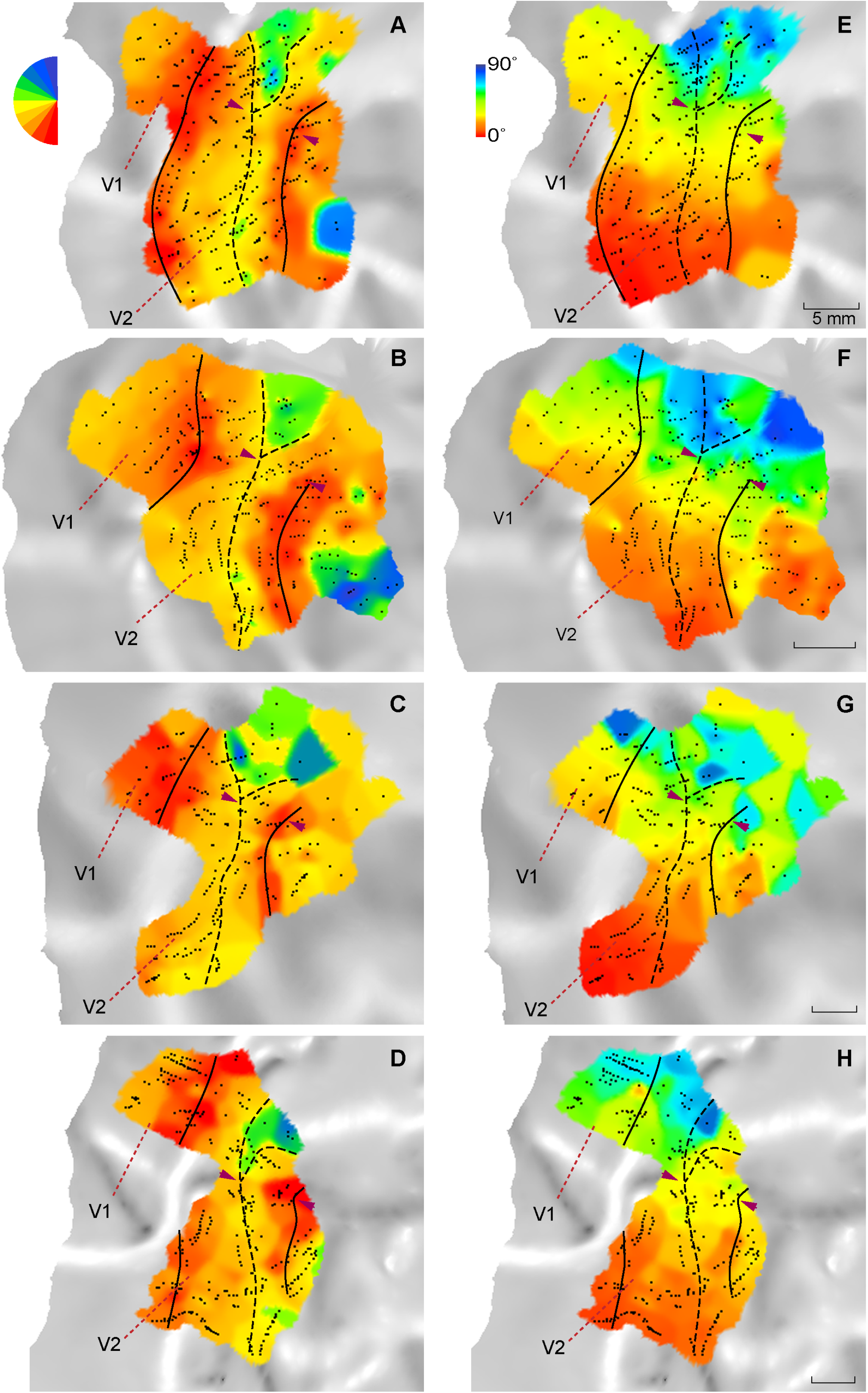
Analysis of visual topography in dorsal extrastriate cortex, shown in unfolded representations of the cortex of the four cases. The receptive field polar angle **(A-D)** and eccentricity **(E-H)** of the recording sites (black points) was interpolated using the method described by Sereno et al. (1994), and colored according to the legends shown adjacent to panels A and E, respectively. In all panels, the black continuous lines were drawn approximately through the middle of the regions of representation of the vertical meridian (red tones, in A-D), and the dashed lines through the regions of representation of the horizontal meridian (yellow/ light green). The pairs of arrow heads point to the region near where the parietooccipital sulcus emerges onto the mesial surface (see text for details).

**Figure 5:**
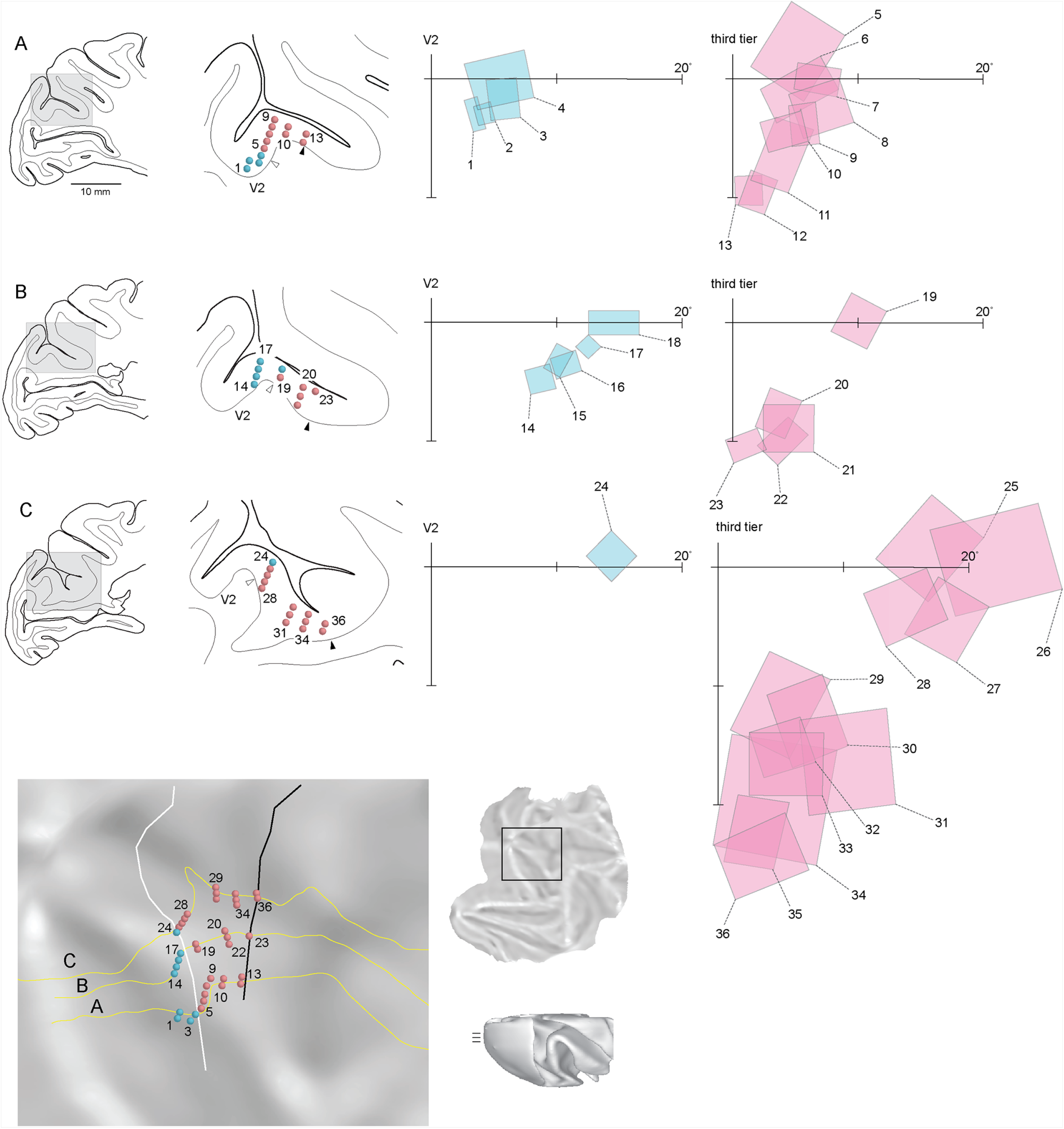
Recording sites and receptive fields obtained from three parasagittal levels **(A-C)** encompassing the annectant gyrus. **Left:** parasagittal sections with recording sites indicated (blue: recording sites assigned to V2; red: recording sites assigned to the area rostral to V2). **Right**: receptive fields recorded in V2 and cortex rostral to V2. **Bottom left**: Cortical layer 4 of the parasagittal sections corresponding to the three levels is indicated by yellow lines, together with the locations of the recording sites. The white and black lines indicate the location of the caudal and rostral borders of the area rostral to V2, estimated as the locations where the receptive field sequences reverted near the horizontal meridian and vertical meridian, respectively. The inserts show the location of the reconstructed region in a flat map prepared with CARET, and the levels of the 3 sections in a dorsal view of the brain.

**Figure 6:**
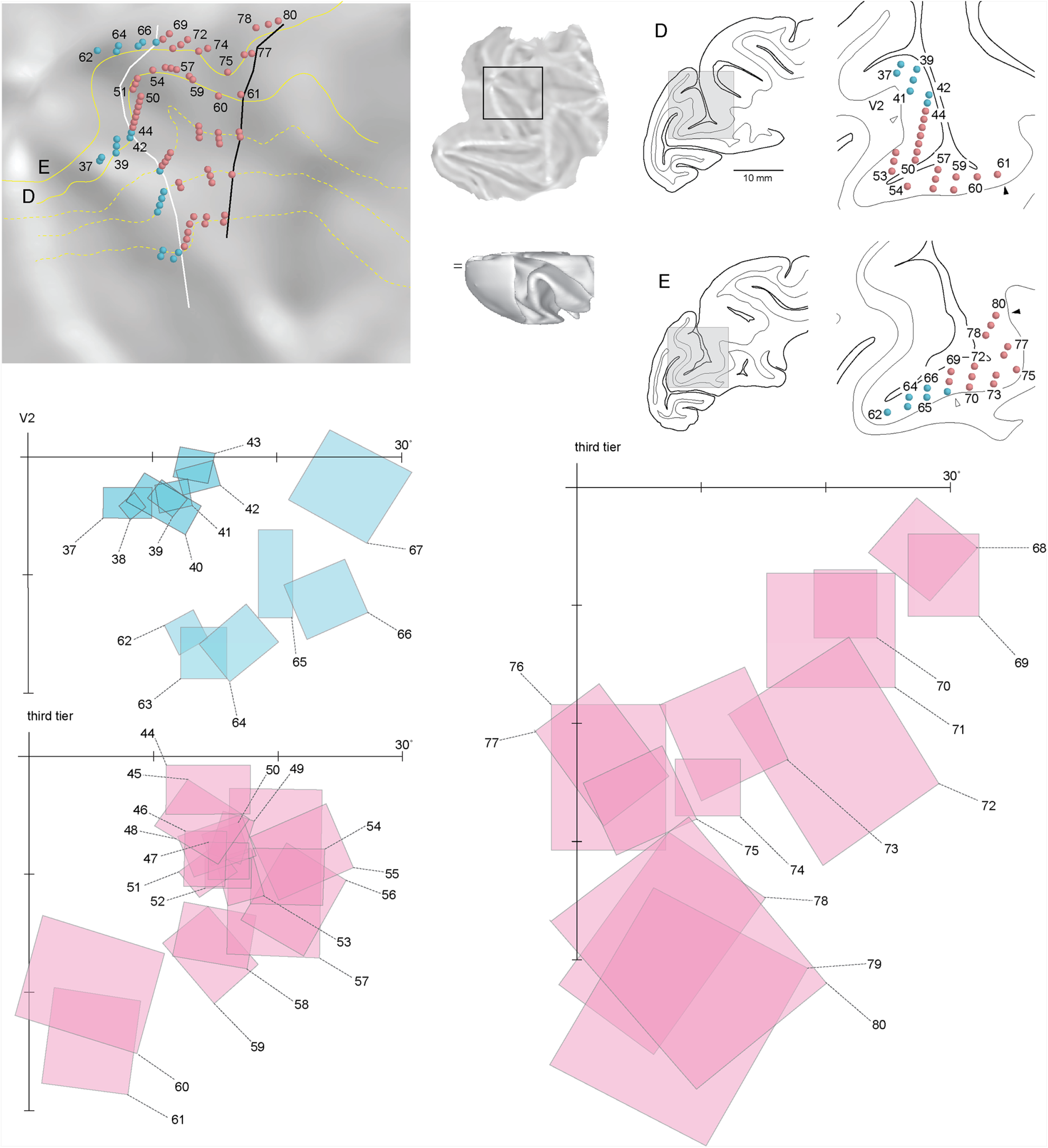
Recording sites and receptive fields obtained from two additional parasagittal levels **(D, E)**, medial to the annectant gyrus. Conventions as in Figure 5.

**Figure 7:**
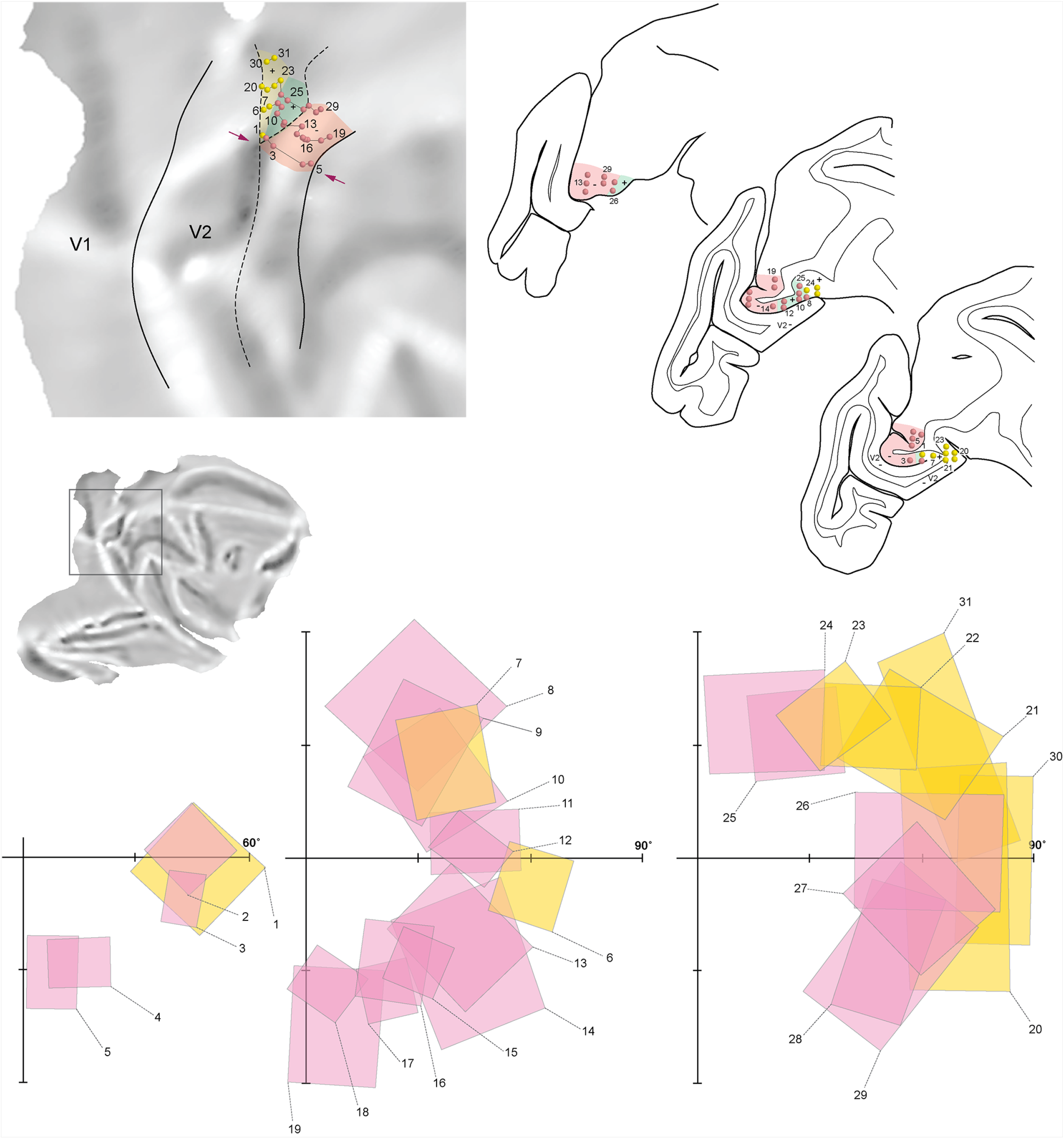
Receptive fields obtained from recording sites at or near the mesial surface. **Top left**: recording sites shown in an unfolded map of the cortex. The solid and dashed lines indicate representations of the lower vertical meridian and horizontal meridian, respectively, and the arrows indicate the region near the emergence of the parieotooccipital sulcus (as shown in Fig. 4). To facilitate visualization of the far peripheral receptive fields in a planar representation, the receptive fields are represented here as rectangles with area proportional to that obtained using appropriate geodesic calculations (Yu and Rosa 2010). Recording sites from different sections were joined into sequences (1-5, 6-19, 20-29 and 30-31) to illustrate the receptive field transitions rostral to V2 (the corresponding receptive fields are shown in the bottom part of the figure). The red recording sites and receptive fields were assigned to third tier cortex (area PO/V6 according to most studies to date), and yellow indicates another representation of the upper peripheral quadrant (POm), which inserts between V2 and PO/V6. **Top right**: location of the recording sites in parasagittal sections.

### V2

As shown in panels A-D of Figure 4, the caudal and rostral borders of dorsal V2 were evident in each animal, coinciding with representations of the vertical (red tones, and continuous lines to the left of each map) and horizontal (yellow tones, and dashed lines) meridians. The receptive field eccentricity in the explored region of V2 (Fig. 4E-H) increased from approximately 5° (red) to 60° or more (dark blue) from lateral to medial. These observations conform well to the findings of earlier studies (e.g. Gattass et al., 1981).

### V3d

Receptive fields obtained in recording sites immediately rostral to V2 in the lunate sulcus and annectant gyrus were also located in the lower visual field, and exhibited a topography that was consistent across animals (Fig. 4A-D). Historically, receptive fields in this region have been attributed to V3d, and the present data conform with the expected visual topography (Van Essen and Zeki, 1978; Gattass et al., 1988). As progressively more anterior sites were sampled in this region, the receptive fields moved from the vicinity of the horizontal meridian (represented at the V2 border, dashed lines) towards the lower vertical meridian (red tones and right continuous lines in Fig. 4; receptive field sequences A-E in Figs. 5 and 6). More importantly, consideration of recording sites arranged from lateral (e.g. sequence A, Fig. 5) to medial (e.g. sequence E, Fig. 6) revealed a monotonic increase in receptive field eccentricity, without any reversals that would indicate the presence of two distinct lower quadrant representations. The width of the putative V3d (3-5 mm) that mirrored the pattern of representation of eccentricity found in V2 was approximately half that of V2. In three cases, small islands of cortex containing receptive fields centered just above the estimated horizontal meridian were observed within the putative limits of V3d, near the lateral limit of the recording grid (light green tones near the bottom of the maps in Fig. 4A, B, D). However, our recording grid did not include more lateral regions where a more extensive representation of the upper quadrant is reported to exist adjacent to V2 (Zhu and Vanduffel, 2018). Receptive field 5 in Figure 5A exemplifies the upper quadrant invasion in this region.

### Cortex medial to V3d

The representation of the horizontal meridian formed the V2/ V3d border until a point near where the rostral bank of the parietooccipital sulcus reaches the midline. Beyond this region (indicated by arrowheads in the Fig. 4 panels) the representation of the horizontal meridian (dashed lines) bifurcated, with a posterior branch that formed the rostral border of V2, and an anterior branch that formed the posterior border of V3d. The region of cortex between these two branches, which occupies parts of the upper bank of the parietooccipital medial sulcus and of the mesial surface, contained a representation of the far periphery of the upper visual quadrant (green and blue tones in Fig. 4A-D; receptive fields 2, 6-12, 20-25 and 31 in Fig. 7). This region has been traditionally assigned to PO/V6, usually being combined with a lower quadrant representation in the parietooccipital sulcus to form a complete representation of peripheral vision (Fig. 1; Colby et al., 1988; Neuenschwander et al., 1994; Gamberini et al., 2015). However, as illustrated in Figures 4 and 7, this lower quadrant representation can be seen as a straightforward continuation of V3d. The eccentricity of the receptive fields recorded in V3d near the lip of the parietooccipital sulcus (arrowheads in Fig. 4; see also Fig. 6 sequence E, and Fig. 7 sequence 1-5) was between 30° and 40°, in agreement with the findings of Gattass et al. (1988). However, extending V3d to encompass the midline cortex increased the range of represented eccentricities to at least 60° (receptive field centers; the complete contralateral hemifield if one takes into consideration the receptive field extent). Accordingly, a parsimonious interpretation of the data suggests that the lower quadrant representation (V3d) extends to the border of the peripheral upper quadrant representation that is usually assigned to PO/V6.

The visual topography of the region encompassed between the two branches of horizontal meridian representation indicated that this cortex contains not one but two mirror-symmetrical representations of the upper quadrant periphery. For example, in Figure 7, receptive field sequences starting at the V2 border first moved towards the upper quadrant vertical meridian (receptive fields 6-7, and 20-23), and then reverted towards the horizontal meridian (receptive fields 8-12, and 24-26) before continuing into a V3d-like lower quadrant representation (receptive fields 13-19, and 27-29). Whereas the second representation formed a natural continuation of the representation of the far peripheral visual field in V3d (green sector in Fig. 7, top), the first representation (yellow recording sites) is likely to correspond to the parietooccipital medial area (POm) described by Neuenschwander et al. (1994) in the *Cebus* monkey and by Rosa and Schmid (1995) in the marmoset monkey.

### Receptive field size

Figure 8 compares the receptive field sizes observed in dorsal V2, V3d (here defined operationally as only containing receptive fields centered in the lower quadrant or on the horizontal meridian), and in the cortex between these areas that represented the upper quadrant. With few exceptions (described above), the latter were located in the representation of the far periphery of the visual field. These results confirm that the receptive fields of V3d multiunit clusters are on average larger than those of V2 (Gattass et al., 1981; 1988), but also indicate that the upper quadrant receptive fields recorded in the cortex interposed between these areas cannot be distinguished from those of V3d. This result argues against receptive field size being a criterion that could justify the distinction between V3d and PO/V6.

**Figure 8:**
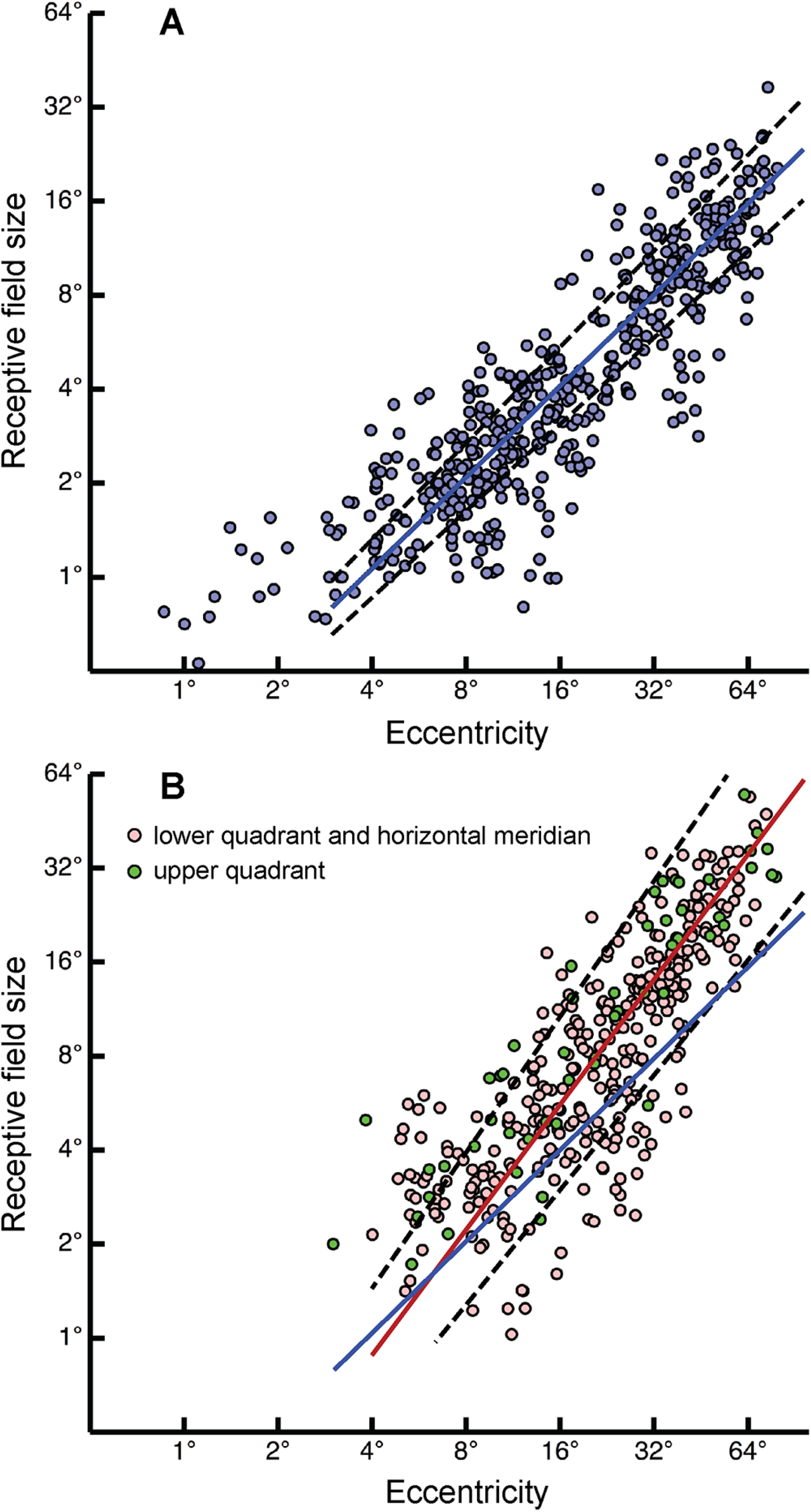
Receptive field size as a function of eccentricity in dorsal V2 **(A)**, and in the cortex immediately rostral to V2, including the territory usually assigned to V3d and PO/V6 **(B)**. Each graph shows receptive fields from 4 animals. The best-fitting power functions, calculated using a model II (principal axis) regression are illustrated for V2 (blue line) and for lower quadrant V3d/PO/V6 (red line). Receptive fields centered on the horizontal meridian were included in this calculation. Confidence intervals (95%) are indicated by dashed lines. The slopes of the functions calculated for lower quadrant V2 and V3d/PO/V6 are clearly different (P< 0.001, permutation test). The number of receptive fields centered on the upper quadrant was much smaller, and non-uniformly distributed across eccentricities, preventing robust statistical comparison. However, the corresponding points (green) overlap extensively with those representing receptive fields in the lower quadrant and horizontal meridian (red). In B, receptive fields assigned to area POm are not included.

## Discussion

We tested the hypothesis that the cortex rostral to V2 in the macaque dorsomedial cortex contains two subdivisions, V3d and V6 (or PO) according to current designations, which form distinct visuotopic maps. This model, which is widely recognized in the current literature, was not supported. Instead, we found that the territory usually assigned to V3d and PO/V6 encompasses a single representation of the lower quadrant, which extends to the temporal limit of the field of vision (Fig. 9). This representation runs as a strip of cortex parallel to V2 for much of its extent, with a shared border formed by neurons that represent the horizontal meridian. However, as it emerges from the rostral bank of the parietooccipital sulcus into the midline, the lower quadrant representation becomes separated from V2 by a region of cortex that represents the periphery of the upper quadrant. At this point, the peripheral visual field is represented according to a simple map that encompasses both quadrants (Fig. 9).

**Figure 9:**
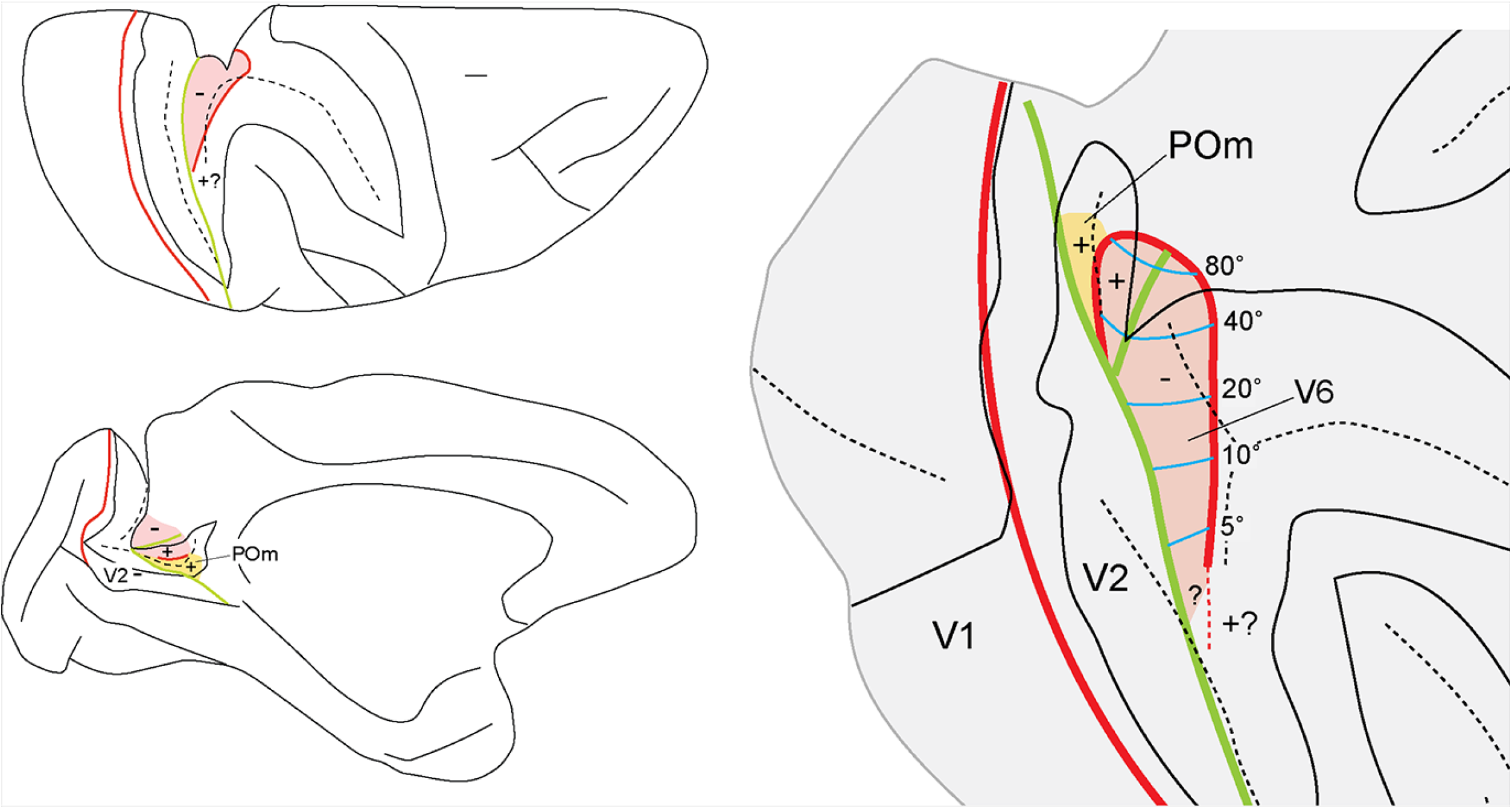
Summary view of the organization of dorsomedial third tier cortex according to present results. **Left**: schematic view of the macaque brain with the sulci partially opened, showing the location of the areas V6 (pink) and POm (yellow). The – and + signs indicate representations of the lower and upper quadrants, respectively, red lines indicate representations of the vertical meridian, and green lines indicate representations of the horizontal meridian. **Right**: unfolded representation of the dorsomedial cortex. Areas V6 and POm are shown with the same conventions. Isoeccentricity lines are shown in blue. The question marks near the lateral end of V6 indicate the likelihood that a representation of the central part of the upper quadrant in this location, similar to that found in New World monkeys, which could complement the upper quadrant representation in V6 (see also Galletti et al. 1999; Rosa and Tweedale 2001).

Most of the representation of the lower quadrant uncovered by our data has been traditionally regarded as part of V3d (Ungerleider and Desimone, 1986; Gattass et al., 1988), whereas the midline region where both the upper and the lower quadrant periphery are represented has been assigned to either PO (Colby et al., 1988) or V6 (Galletti et al., 1999). Comparison with published results strongly indicates that the revised area we propose overlaps well with these subdivisions (Fig. 10). However, our results suggest that V3d (at least, the part located medial to the annectant gyrus) and PO/V6 form a single topographic map, without reversals or re-representation at the putative border. We also found no evidence for an additional area (PIP) separateing V3d from PO/V6, insofar as the receptive field topography is concerned. Our results therefore support the view that V3d, PO and V6 are best regarded as designations which have been applied to parts of a same area. This interpretation is supported by well-documented anatomical similarities between V3d, V6 and PO, reviewed elsewhere (e.g. Rosa et al. 2005, 2009; Angelucci and Rosa, 2015). Each of these putative areas has been described as being heavily myelinated relative to adjacent cortex (see also Fig. 2), and to receive a dense projection from layer 4b of V1, which makes them different from ventral and lateral parts of the third tier complex (Colby et al., 1988; Felleman et al., 1997; Galletti et al., 2001). Together, these results argue against the notion that most of the cortex adjacent to V2 is formed by a single area which includes upper and lower quadrant representations located segregated to ventral and dorsal extrastriate cortex, respectively (e.g., Gattass et al., 1988; Lyon and Connolly, 2012).

**Figure 10:**
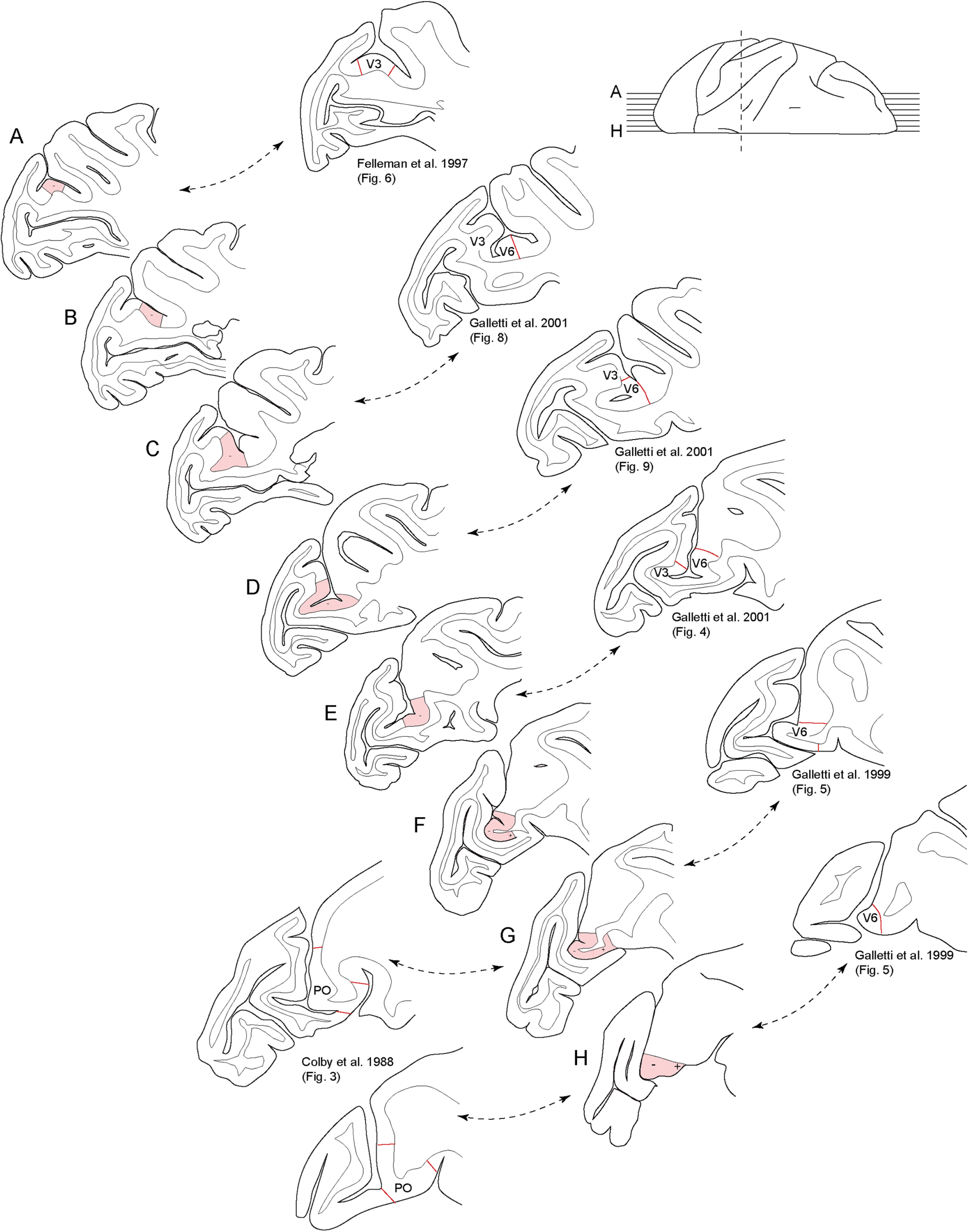
**A-H**: parasagittal sections showing the extent of the redefined area V6 according to the present results (pink). For comparison, the left and right columns show drawings of sections from previous publications showing the extents of areas V3 (Felleman and Van Essen 1997), V6 (Galletti et al. 1999, 2001) and PO (Colby et al. 1988). The presently redefined V6 overlaps extensively with the cortex occupied by these areas recognized by previous studies.

### Nomenclature, and comparison with previous models

Our findings raise the issue of what is the most appropriate designation for the dorsomedial third tier area. On balance, this area most closely resembles area V6 as proposed by Galletti et al. (1999, 2001). Like V6, the area encompasses a strip-like central and peripheral lower quadrant representation, as well as a peripheral upper quadrant representation in the mesial surface and parietooccipital sulcus. Thus, in order to avoid multiplication of names, we favor retaining the designation V6, with the provision that this area is not separated from V2 by a strip of V3d, as proposed previously (e.g. Gamberini et al., 2015).

Another viable option would be to retain the name V3d, which has historical precedence (Ungerleider and Desimone, 1986), and has anatomical characteristics similar to those of the area we mapped (Burkhalter et al., 1986). Although there is some potential for confusion (given that V3d is traditionally seen as only forming a lower quadrant representation), this would mitigate to some extent the reasons why V3d has been regarded as an “improbable area” (Zeki, 2003).

Finally, we regard the designation PO as the least satisfactory. As discussed by Galletti et al. (2005), this area, as defined by Colby et al. (1988), seems to extend further dorsally than V6 does, thereby encompassing parts of the parietooccipital sulcus which are now known to belong to a functionally different area (V6Av; Passarelli et al., 2011).

Regardless of the final verdict on nomenclature, our results align with the fact that there are well known anatomical and functional differences between the dorsal and ventral components of the third tier cortex (the latter encompassing the “ventral subdivision of V3” [V3v], or ventral posterior area [VP]; Burkhalter et al., 1986). In view of the above, the designation V3 could be assigned to this area, which forms most of the border of V2 (including the ventral surface, caudal prelunate gyrus, and possibly lateral parts of the lateral lunate sulcus; Rosa et al., 2000; Rosa and Tweedale, 2001). The designation V3 would reflect the characteristics of the homonymous area in other mammals, which is g involved in object shape analysis (Manger and Rosa 2005).

### Comparison with New World monkeys

Many of the features of the revised V6 resemble those of the DM, as defined in the marmoset monkey (Rosa and Schmid 1995). Like the revised macaque V6, DM also shows a continuous complete representation of the lower quadrant adjacent to V2, adjoined by a representation of the peripheral upper quadrant in the midline cortex (Fig. 9; Rosa and Tweedale, 2001). DM also shares the dense myelination of V6, and its specific, dense input from layer 4b of V1 (which is known as layer IIIc in the nomenclature of Hassler, 1996, favored in many studies of New World monkeys; see Elston and Rosa, 1997 for discussion). Moreover, the connections of DM appear identical to those of V3d and PO/V6, if one allows for the interpretation that these are central and peripheral representations of a same area (Rosa et al., 2009). Because the grid of electrode penetrations did not extend far into the banks of the lunate sulcus, our data do not establish the full lateral extent of the redefined macaque V6. Whereas the traditional model proposes that its lateral part (V3d) extends adjacent to V2 all the way to the lateral end of the lunate sulcus (Gattass et al. 1988), more recent studies have questioned this, proposing instead that part of the cortex rostral to V2 in this sulcus represents the central upper quadrant (Zhu and Vanduffel, 2018; meta-analysis by Angelucci and Rosa, 2015). Although we did observe a few receptive fields centered in the upper quadrant in the expected location, the sample was insufficient to ascertain the extent of this representation. This is unfortunate, because our data leaves open the important question of whether receptive fields recorded in the lunate sulcus complement the upper quadrant representation we identified in the midline, as observed in area DM of marmoset monkeys (Rosa et al., 2005; 2009; Angelucci and Rosa, 2015). Thus, although the present results indicate that the medial part of V3d and PO/V6 are best considered as parts of the same area, establishing the extent of the similarity of this area to the New World monkey DM will require further work. Earlier studies (e.g., Beck and Kaas, 1999) proposed that the designation DM would also be appropriate for this part of the macaque cortex, and we suggest that V6 and DM could be seen as equivalent designations for the homologous areas in the macaque and marmoset.

### Area POm

Our data also revealed an additional representation of the peripheral upper quadrant in the parietooccipital medial sulcus of the macaque, which inserts between the peripheral representations of V2 and V6. The region has also been observed in electrophysiological studies of marmosets and capuchins (Neuenschwander et al., 1994; Rosa and Schmid, 1995), which suggested the designation area POm; similar to these studies we found that POm could be distinguished from V6 by its lighter myelination (cortex rostral to PO/V6 in the parietooccipital sulcus, in Fig. 2). However, our recordings were insufficient to establish the full extent of POm. POm appears to overlap with the scene-selective retrosplenial region identified by Nasr et al. (2011) and with the medial visual region (Vis) reported by Passarelli et al. (2018) in the macaque. It may also correspond to the recently described human brain area V2A (Elshout et al., 2018).

### Conclusions

Our findings point to a simpler model of the organization of the dorsomedial cortex anterior to area V2 in macaques, which consolidates subdivisions proposed by different authors into a single area. We propose retaining the designation V6 for this area, while acknowledging the likelihood that it is homologous to area DM in New World monkeys. This model provides a parsimonious account of previous anatomical and electrophysiological observations, including differences between the dorsal and ventral components of the third tier complex (Burkhalter et al., 1986; Beck and Kaas, 1999; Rosa et al., 2005; Jeffs et al., 2015). Knowing the extent of its application to the human brain will require high-field imaging studies that include stimulation of the far periphery of the visual field, preferably combined with appropriate structural and functional connectivity experiments.

## Acknowledgements

The authors acknowledge the contributions of Rowan Tweedale during many phases of this project, including participation in experiments, and critical reading of earlier versions of the manuscript. We also thank Janssen-Cilag for the provision of sufentanil citrate.

## Author Contributions

KH, SB and MGPR designed research; KH, SB, HHY and MGPR performed experiments; KH, SB, TAC, HHY, OA, JMC and MGPR analyzed the data; KH, SB and MGPR wrote the paper.

